# Multi-omics analysis of the potential of MACC1 as a biomarker and therapeutic target for colonic adenocarcinoma

**DOI:** 10.64898/2026.03.21.713326

**Authors:** Zhenkun Chen, Chun Zheng, Yi Zhang, Xulin Peng, Yang Lu, Jie Zhang, Peng Sun

## Abstract

Colonic adenocarcinoma (COAD) is a major cause of cancer-related mortality worldwide. Various tumors are linked to metastasis-associated in colon cancer 1 (MACC1). This study aimed to analyze public datasets to examine MACC1 expression, signaling pathways, copy number variations, and associations with immune cell subsets in COAD employing bioinformatics. MACC1 expression was elevated in COAD, especially in Wnt signaling and chromatin modifier pathways. Analysis of somatic copy number alterations in The Cancer Genome Atlas-COAD dataset revealed a link between MACC1 and DNA damage repair. MACC1 also showed a negative correlation with genes involved in immune cell infiltration in patients with COAD, including cluster of differentiation (CD)8+ T cells, activated dendritic cells, CD8 T cells, and cytotoxic cells. Collectively, these findings suggest MACC1 as a potential prognostic biomarker and therapeutic target for COAD.

## Background

The development and progression of colonic adenocarcinoma (COAD) involve multifaceted genetic and molecular alterations. Among various genes implicated in this malignancy, metastasis-associated in colon cancer 1 (MACC1), which was initially identified in 2009, has emerged as a pivotal gene. Extensive evidence validates MACC1 as an independent prognostic biomarker and a critical driver of metastasis in colon cancer, particularly COAD, highlighting its clinical and biological significance [1]. This multifactorial involvement further positions it as a master regulator of tumor aggressiveness and a promising therapeutic target.

At the molecular level, MACC1 exerts its oncogenic effects primarily by activating the hepatocyte growth factor (HGF)/cellular mesenchymal–epithelial transition (MET) factor (c-MET) signaling axis, which drives tumor growth, angiogenesis, and metastasis. Reportedly, MACC1 binds directly to the MET promoter, boosting c-MET expression and downstream malignant signaling, driving malignant phenotypes in COAD cells [2,3]. MACC1’s mechanisms extend beyond the classical HGF/c-MET pathway, encompassing a broader spectrum of regulatory layers, to include transcriptional control, epigenetic modifications, and post-transcriptional regulation. Epigenetic regulations are also critically implicated in the regulation of MACC1 expression and function [4,5]. Beyond transcriptional and epigenetic regulations, MACC1 reprograms cancer cell metabolism, which is critical to tumor progression, via metabolic reprogramming, including promoting glycolysis via glucose transporter type 4 membrane translocation in colorectal cancer cells [6]. This metabolic shift meets the energetic and biosynthetic demands of proliferating tumors, contributing to MACC1 overexpression-associated aggressive phenotype. Emerging evidence correlates MACC1 with immune evasion mechanisms via immune cell infiltration and immune checkpoint molecules [4]. This immunomodulatory function of MACC1 complexifies its role in tumor biology, highlighting its potential for combined immunotherapies.

The clinical relevance of MACC1 stems from its strong links to poor prognosis, advanced tumor stage, lymph node involvement, and distant metastasis across cancers, including COAD [4,7]. Beyond its role as a prognostic biomarker, MACC1 can act as a predictive marker for therapeutic response [8]. Additionally, combination therapies inhibiting MACC1-dependent drug resistance mechanisms, such as ATP-Binding Cassette subfamily B member 1-mediated chemoresistance, have shown improved outcomes in colorectal cancer cells [9]. Collectively, these therapeutic insights emphasize the translational potential of MACC1-targeted interventions.

While molecular classification aids COAD diagnosis, no definitive early biomarker has been identified for this malignancy. Consequently, molecular subtyping should be regarded as complementary to, rather than a full replacement for, traditional histomorphological classification. Integrating molecular classifications into prognosis refines risk stratification and may improve clinical management for specific patient subsets [10,11].

This study examined MACC1’s multifaceted roles in COAD employing bioinformatics and public datasets to assess its potential as a novel predictive or therapeutic marker in COAD. Expression profiling of MACC1 across tumor stages and histological subtypes showed its upregulation at both transcriptional and protein levels, which correlated with unfavorable outcomes. Furthermore, MACC1 showed a relationship with immune cell infiltration in the tumor microenvironment. Somatic copy number alterations (SCNAs) in MACC1-enriched COAD samples also highlighted pathways such as DNA damage repair and immune-metabolic reprogramming, linking MACC1 to therapy responsiveness.

## Results

### Screening process for *MACC1* relates to cell growth in COAD

Through an intersectional analysis of gene sets from Gene Expression Profiling Interactive Analysis (GEPIA2), we identified key genes with both statistical robustness and biological relevance. These genes likely contributed to core disease mechanisms, including signal transduction, cell cycle regulation, and apoptosis. One set comprised 390 genes significantly associated with COAD prognosis (p < 0.05), whereas the other included 5,222 genes differentially expressed between COAD and normal tissues (p < 0.05). Their intersection yielded 109 genes, visualized in a Venn diagram created using the online Xiantao tool (Fig. 1A). Subsequent Gene Ontology (GO) and Kyoto Encyclopedia of Genes and Genomes (KEGG) functional analyses revealed enriched pathways and biological processes (Fig. 1B). From the GO results, genes linked to “regulation of Wnt signaling pathway” were further intersected with cancer-related genes from “regulation of non-canonical Wnt signaling pathway,” highlighting MACC1 as a key candidate. Given its established role as an oncogene, involvement in multiple signaling pathways, and documented alterations in colorectal cancer (particularly COAD), MACC1 was selected as the focus of this study.

**Fig. 1.**
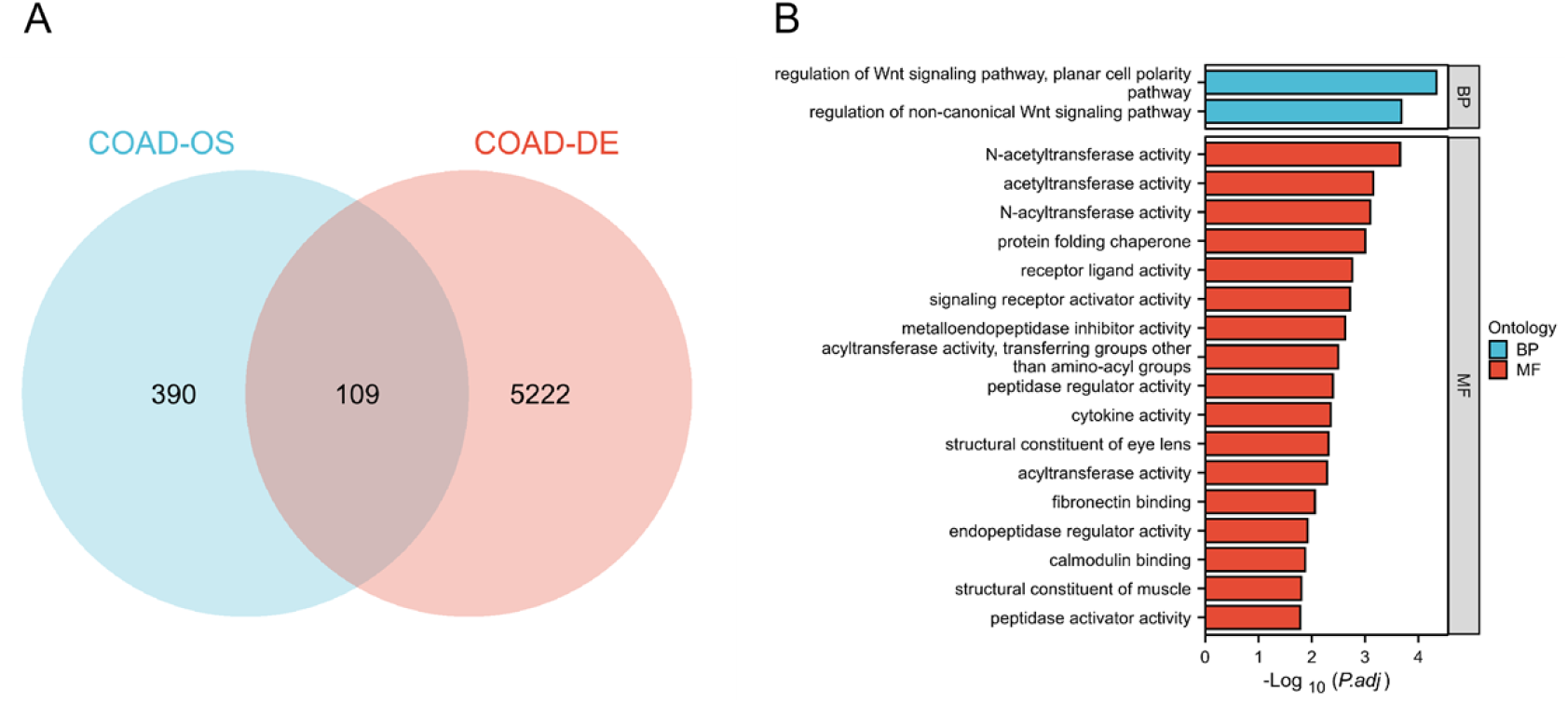
Screening process for target genes and MACC1 related to the Wnt-signal pathway in COAD. (**A**) Venn diagram for genes associated with survival and differential expression. (**B**) GO and KEGG enrichment analyses showed different functions of 105 intersecting genes.

### MACC1 transcription level is upregulated in COAD and associated with poor prognosis

We first analyzed MACC1 transcriptional expression across 33 cancer types, including COAD (Fig. 2A). In COAD, MACC1 mRNA levels were significantly higher in 480 primary tumor samples than in 41 normal tissues (Fig. 2B), and elevated expression persisted across advancing tumor stages (Fig. 2C) and histological subtypes (Fig. 2D). This MACC1 upregulation was correlated with disease progression and tumor severity. Furthermore, increased MACC1 transcription correlated with poorer survival in patients with COAD (Fig. 2E).

**Fig. 2.**
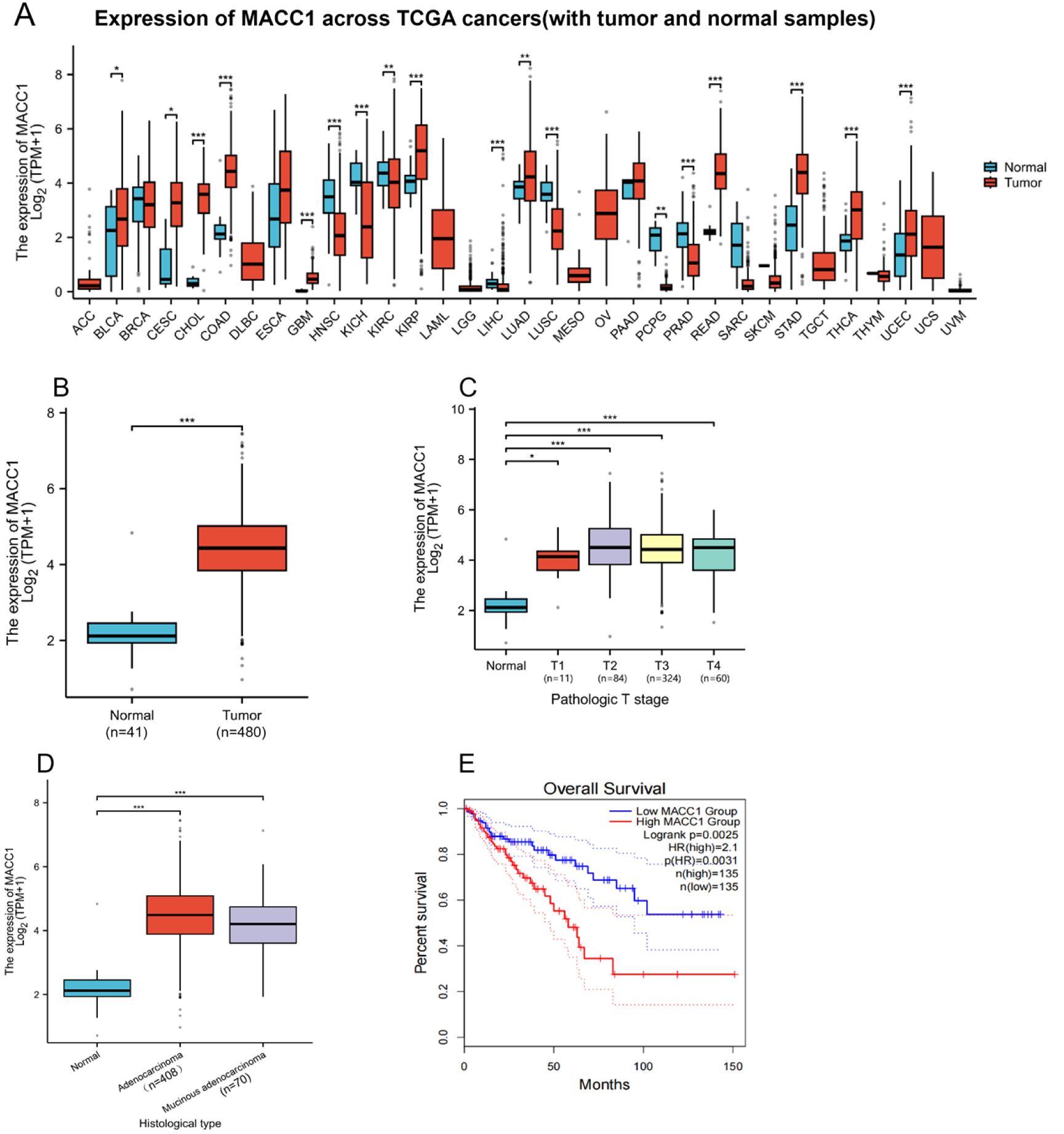
MACC1 is upregulated in COAD patients and associated with poor prognosis. **(A)** MACC1 expression in pan-cancer. Blue represents normal, red indicates a tumor. **(B)** MACC1 expression between normal and primary tumor tissues. (**C**) MACC1 expression in different cancer stages. (**D**) MACC1 expression in different histological subtypes. (E) Overall survival rate of low and high MACC1 expression groups. A total of 270 tumor samples from GTEx were analyzed. Log-rank *p* = 0.0025. Group cutoff: median. Hazard ratio (HR) = 2.1, *p* = 0.0031. HR was calculated based on the Cox PH model. 95% confidence interval is indicated as a dotted line. \**p* < 0.05; \*\**p* < 0.01; \*\*\**p* < 0.001.

### MACC1 protein expression is upregulated in COAD

MACC1 protein levels were evaluated using data from the Clinical Proteomic Tumor Analysis Consortium (CPTAC) and The University of ALabama at Birmingham CANcer data (UALCAN) portals. Analysis of 100 normal and 97 COAD samples revealed that MACC1 protein levels positively correlated with COAD (Fig. 3A), with its significant upregulation in early-stage tumors and non-mucinous histological subtypes (Fig. 3B, C). Immunohistochemistry (IHC) images from the Human Protein Atlas (HPA) corroborated these findings. Normal colonic mucosa showed low or undetectable MACC1 expression, whereas tumor tissues exhibited strong immunostaining (Fig. 3D; representative IHC results). These data indicate elevated MACC1 expression during early stages in COAD progression, notably at stage 1 and in non-mucinous subtypes. Given its clarity and practicality, IHC holds promise as a tool for early tumor detection.

**Fig. 3.**
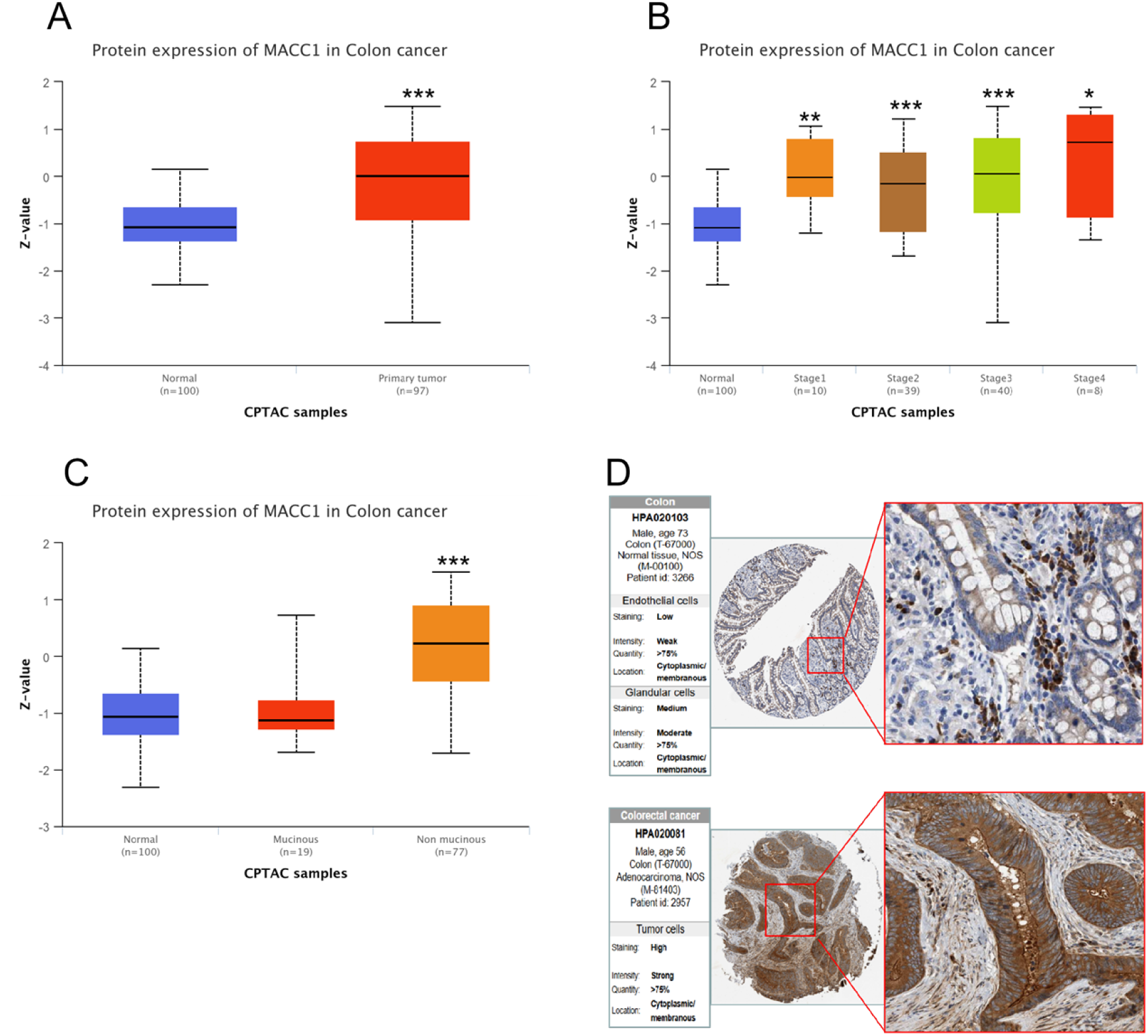
MACC1 protein expression is upregulated in COAD patients. (A) MACC1 protein expression increases significantly in COAD samples. Z-values represent standard deviations from the median across samples. MACC1 protein expression in different COAD (B) stages, (C) histological subtypes, (D) IHC image of MACC1 in normal and COAD tissues. \**p* < 0.05; \*\**p* < 0.01; \*\*\**p* < 0.001.

### Two MACC1-related signaling pathways are involved in COAD

To explore MACC1’s role in COAD pathogenesis, we analyzed its links to signaling pathways using the CPTAC-UCEC dataset (including 94 COAD cases), classifying samples as “pathway altered” or “others” based on the alteration status of specific pathways. MACC1 protein levels were compared between these groups and 100 normal adjacent tissues. Notably, MACC1 levels strongly correlated with multiple signaling pathways, including Hippo, Wnt, p53/Rb, mechanistic target of rapamycin (mTOR), receptor tyrosine kinase (RTK), myelocytomatosis oncogene (MYC)/myelocytomatosis oncogene neuroblastoma (MYCN), SWItch (SWI)–Sucrose Non-Fermentable (SNF), chromatin modifiers, and nuclear factor erythroid 2-related factor 2 (NRF2) pathways (Fig. 4). Importantly, MACC1 showed pronounced correlations with altered Wnt and chromatin modifier pathways (red box).

**Fig. 4.**
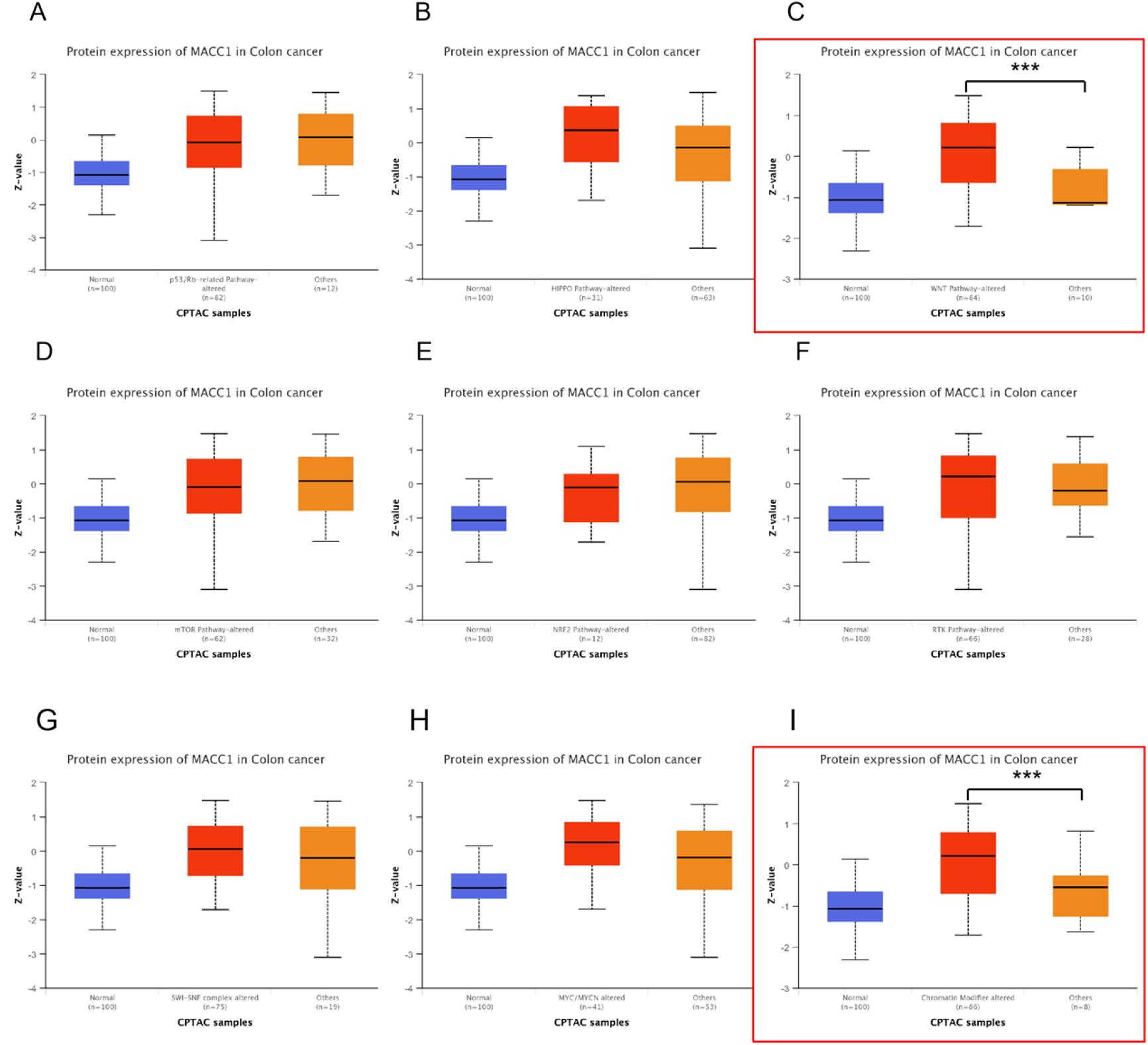
Alteration in Wnt- and chromatin-modifiers related pathways associated with MACC1 protein expression. UALCAN analysis of MACC1 proteomic expression based on Hippo, Wnt, p53/Rb, mTOR, RTK, MYC/MYCN, SWI/SNF complex, chromatin modifier, and NRF2 status. MACC1 protein expression was determined using mass spectrometry. Z-values represent standard deviations from the median across COAD samples. A total of 944 CPTAC-COAD cases were used. \**p* < 0.05; \*\**p* < 0.01; \*\*\**p* < 0.001.

### MACC1 expression correlated with immune cell infiltration in COAD

Single-sample gene set enrichment analysis (ssGSEA) and Spearman correlation results revealed the relationship between MACC1 expression and immune cell infiltration levels in COAD (Fig. 5A). Notable negative correlations were found between MACC1 expression and the number of activated dendritic cells (aDCs) (*R* = −0.336, *P* < 0.001; Fig. 5B), cytotoxic cells (*R* = −0.364, *P* < 0.001; Fig. 5D), and cluster of differentiation (CD)8 T cells (*R* = −0.187, *P* < 0.001; Fig. 5C); MACC1 expression positively correlated with T central memory (Tcm) cells (*R* = 0.396, *P* < 0.001; Fig. 5E) and T helper cell (*R* = 0.208, *P* < 0.001; Fig.5F).

**Fig. 5.**
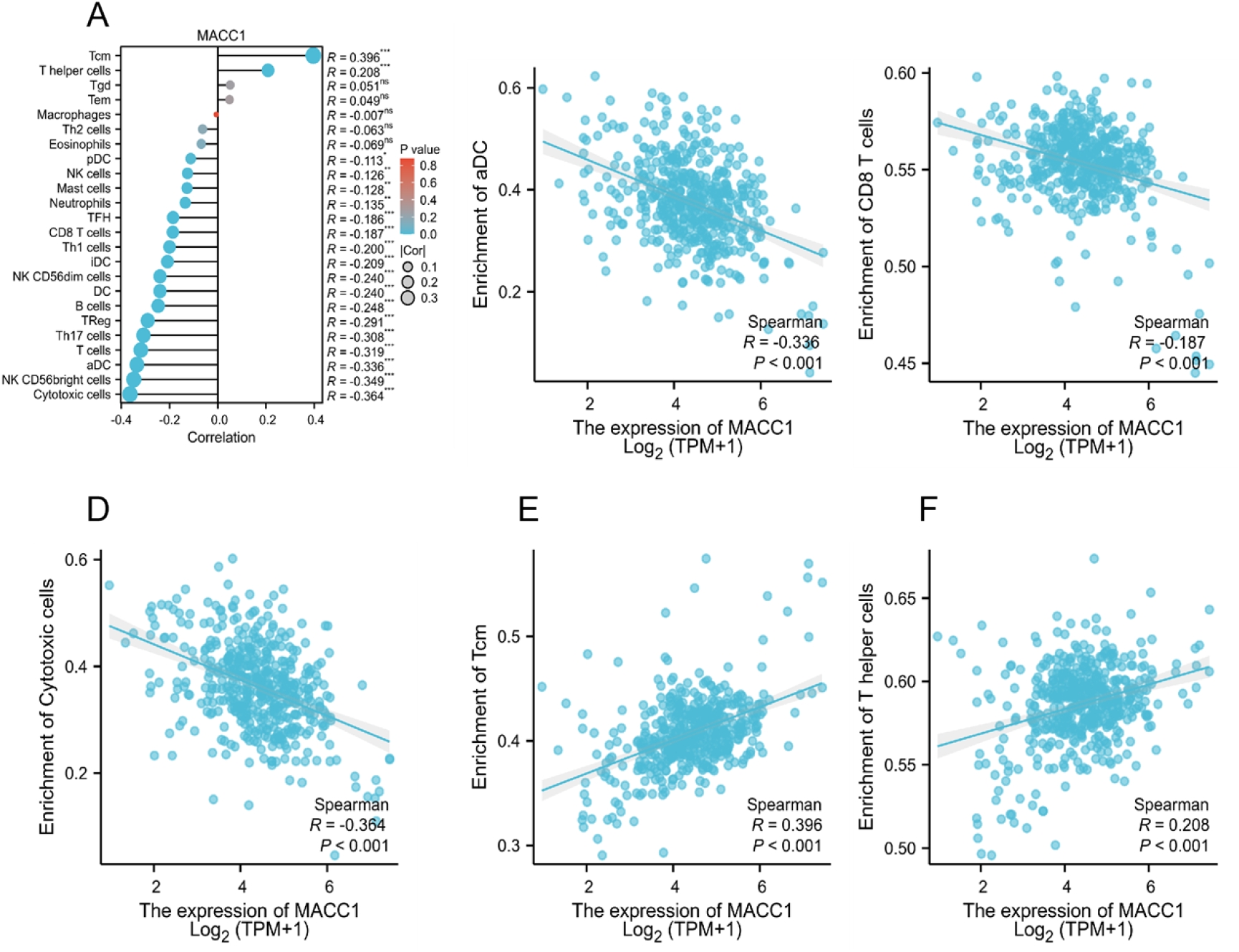
Relationship between MACC1 expression in the tumor microenvironment of COAD and immune cell infiltration. **A** Relationship between immune cell levels and MACC1 mRNA expression. The relationships between the abundances of (**B**) aDC, (**C**) CD8 T cells, (**D**) cytotoxic cells, (**E**) Tcm, (**F**) T helper cells are shown in scatter plots.

### MACC1 SCNAs relate to signaling pathways in COAD

SCNAs are a common genomic feature in human cancers and drive tumorigenesis by modulating gene expression. Aberrant DNA copy number variations (CNVs) represent underlying molecular mechanisms of various diseases, including cancer and metabolic disorders. In COAD, CNVs have been closely linked to patient prognosis. Using the LinkedOmics platform, we analyzed MACC1 CNV data from The Cancer Genome Atlas (TCGA)-COAD cohort (n = 629) alongside matched RNA-sequencing profiles for correlation analysis. A volcano plot (Fig. 6A) revealed genes with significant positive (dark red dots) or negative correlations (dark green dots) correlations to MACC1-SCNAs. Heatmaps depicted expression patterns of the top correlated genes (Fig. 6B, C). Functional enrichment analysis of up- and down-regulated gene sets identified biological processes tied to MACC1 CNV status (Fig. 6D). Drug sensitivity analyses via GDSC and CTRP databases linked MACC1 levels to pharmacological responses (Fig. 6E). MACC1 levels correlated with sensitivity to various compounds: negative correlations with CEP-710, FK-866, AZD4547, and TW37; positive correlations with BRD-A86708339 and Lapatinib.

**Fig. 6.**
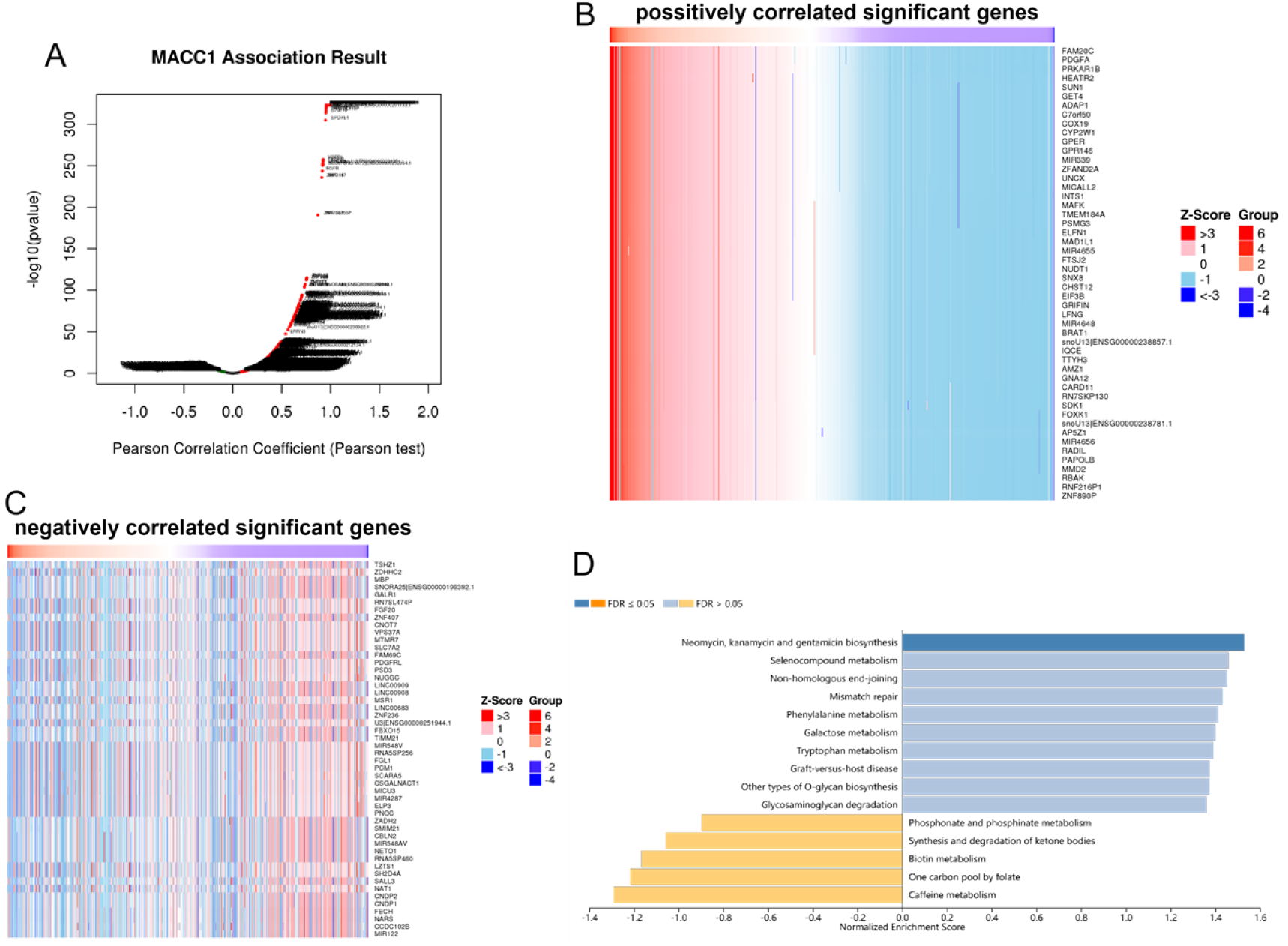
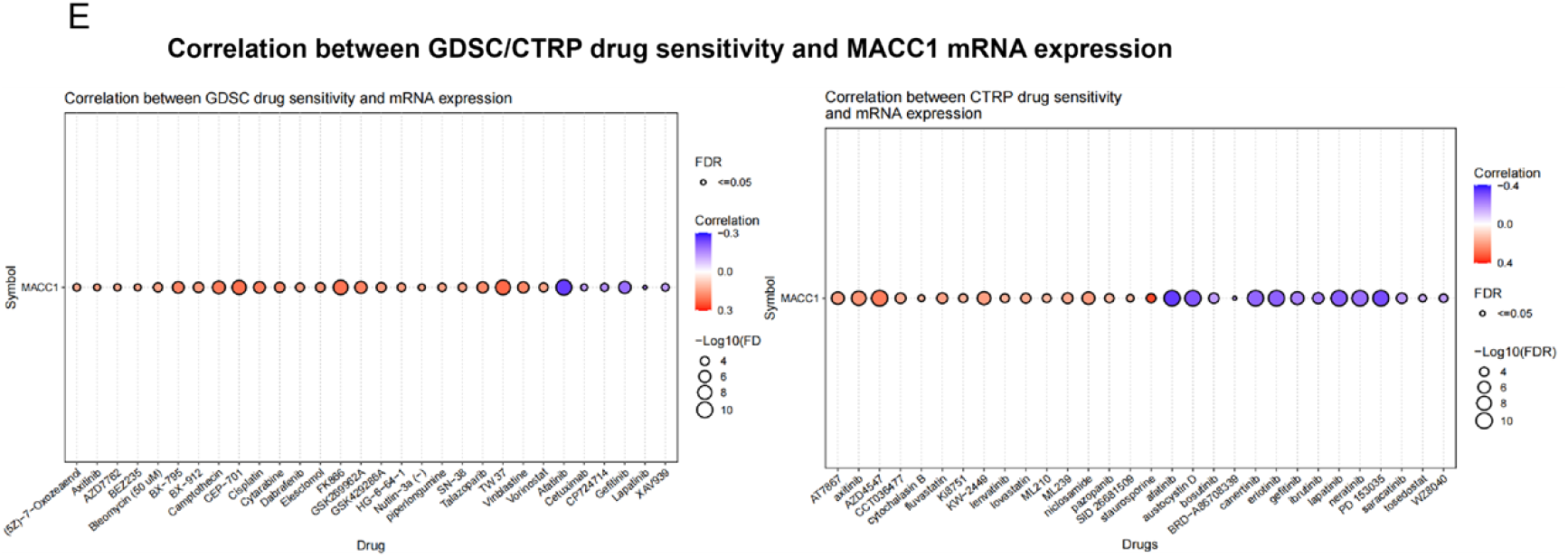
Somatic copy number alteration in MACC1 is related to drug metabolism in COAD. (A) Differential genes from RNA-seq and COAD mutant-related SCNA data for TCGA-COAD differential genes. P-value obtained from Pearson’s correlation test. (B,C) Heatmap of differential genes. (D) Top 15 KEGG pathways for differential genes. (E) Correlation between MACC1 expression and sensitivity of GDSC and CTRP drugs (top 30) in pan-cancer. Orange displays positive correlation; purple represents a negative correlation.

## Discussion

*MACC1*, located on human chromosome 7p21.1, was initially identified as a key driver of metastasis in colon cancer. MACC1 protein features several functional domains, including a critical Src homology 3 (SH3) domain that interacts with downstream signaling molecules to mediate oncogenic processes [1].

In lung adenocarcinoma, MACC1 binds the splicing factor HNRNPH1 via its SH3 domain, regulating aberrant splicing of oncogenes [12]. MACC1 exhibits heterogeneous subcellular localization in the cytoplasm, nucleus, and cell membrane, reflecting its diverse functional roles. Notably, nuclear MACC1 acts as a transcription factor; for instance, it binds directly to the MET gene promoter to enhance transcription [13]. Similarly, MACC1 can bind to the *LGR5* promoter region to promote cancer stem cell characteristics [14]. MACC1 expression undergoes multifaceted and precise regulation. At the transcriptional level, various transcription factors influence its expression. In colon cancer, factors such as c-JUN, hepatocyte nuclear factor 4-gamma, and paired box 6 bind the MACC1 promoter, indicating their role in the transcriptional regulation of MACC1 [15]. The pro-inflammatory cytokine tumor necrosis factor-alpha also induces MACC1 via nuclear factor-κB subunit p65 and c-JUN, promoting tumor growth and metastasis [16].

Epigenetic modifications further drive MACC1 overexpression. In colorectal cancer, reactive oxygen species-induced oxidative stress upregulates MACC1 by altering histone modification states, specifically by upregulating active mark H3K4me3 and downregulating repressive mark H4K20me3. This alleviates epigenetic silencing of MACC1 [5]. Chromosomal instability and SCNA also contribute significantly to MACC1 overexpression in colorectal cancer, which, in turn, is associated with a poor survival prognosis [17].

Non-coding RNAs act as key negative regulators of MACC1 expression. For example, miR-638 targets the MACC1 3’ untranslated region to suppress its expression, thereby inhibiting proliferation, migration, and invasion in gastric cardia adenocarcinoma cells [19]. Conversely, in colorectal cancer, circular RNA circ-0006174 upregulates MACC1 by sponging miR-138-5p, promoting cancer progression [18].

Tumor progression involves multiple, intricate signaling pathways, with MACC1 serving as a key upstream regulator of the HGF/c-MET signaling. MACC1 is well-characterized for its molecular function in transcriptionally activating MET by binding its promoter [13]. In colorectal cancer, MACC1, acting as an upstream regulator of c-MET, enhances c-MET signaling, driving cell proliferation and metastasis following its overexpression [2]. MACC1 also engages in bidirectional regulation with the Wnt/β-catenin pathway, forming a robust oncogenic network. The β-catenin/transcription factor 4 complex, a core effector molecule of the Wnt/β-catenin pathway, transcriptionally activates MACC1 expression [19].

MACC1 also functions as a transcription factor that can bind to the promoter of *LGR5*, a marker of cancer stem cells, to induce cancer stem cell characteristics in colorectal cancer [14]. Additionally, MACC1 promotes epithelial–mesenchymal transition (EMT) by interacting with the EMT regulator Snail family transcriptional repressor 1 (SNAI1), as seen in pancreatic cancer. Notably, EMT is also mediated by Wnt/β-catenin to facilitate metastasis [20]. This strong positive feedback loop between MACC1 and Wnt/β-catenin centrally drives the malignancy in colon cancer. Consequently, targeting this reciprocal interaction with combined blockade strategies may prove more effective in inhibiting colon cancer progression than targeting either pathway alone [14].

Mechanistically, MACC1 acts as a transcription factor that directly binds to and activates SNAI1, a key EMT regulator. This upregulates mesenchymal markers such as fibronectin 1 and downregulates epithelial markers such as E-cadherin, promoting a mesenchymal phenotype in cells [21]. Furthermore, MACC1 expression is significantly correlated with various EMT-related transcription factors, including Snail and Twist, as well as with poor prognosis. Various studies validate this correlation across cancers, including hepatocellular carcinoma and esophageal squamous cell carcinoma [20,22].

MACC1 profoundly shapes the tumor immune microenvironment. High MACC1 levels closely correlate with altered immune cell infiltration. In breast cancer, elevated MACC1 associates positively with CD163, a marker of the M2 protumor phenotype in tumor-associated macrophages, while negatively correlating with cytotoxic CD8+ T cells and CD56+ natural killer cells. This indicates that MACC1 may contribute to the establishment of an immunosuppressive microenvironment, enabling tumor immune evasion [23].

Investigating MACC1 as a molecular switch in COAD malignancy has deepened insights into tumor biology beyond traditional c-MET activation. From an expert perspective, MACC1 operates in a complex, multidimensional regulatory network coordinating transcription, signaling, tumor microenvironment remodeling, and metabolic reprogramming. This integrative role highlights the intricacy of cancer pathogenesis and the importance of holistic approaches in exploring oncogenic drivers.

Balancing diverse research perspectives, MACC1 integrates multiple oncogenic signals to empower tumor cells, driving proliferation, invasion, and metastasis, while simultaneously mediating resistance to chemotherapy and radiotherapy.

Clinically, MACC1 serves as a potent prognostic biomarker and promising liquid biopsy target, advancing precision oncology in colorectal cancer. Its expression strongly correlates with patient outcomes, enabling refined risk stratification and real-time therapeutic monitoring [7,24,25]. This translational potential links molecular research to patient care, supporting tailored treatments and improved survival. However, clinical implementation of MACC1-based diagnostics and therapeutics requires standardized assays, validation in large, diverse cohorts, and workflow integration.

Herein, MACC1 expression was elevated in COAD and correlated with shorter patient survival. Notably, MACC1 upregulation appeared early in tumor progression, accompanied by increased protein levels. IHC analysis from the HPA database further confirmed higher MACC1 protein in COAD tissues versus normal colonic mucosa.

MACC1 expression inversely correlated with CD8+ T cells, which play a critical role in immune responses against infections and tumors. Reduced CD8+ T cell activity aligned with elevated MACC1 levels (Fig. 5), implying a negative association. The underlying mechanisms require further elucidation. Proteomic mass spectrometry linked CDKN2A expression to Wnt signaling and chromatin modifier pathways. This finding further motivates exploration into the possible involvement of MACC1 in tumor progression.

CNVs, including genomic duplications or deletions, are common in COAD. TCGA data show that high-CNV-burden COAD cases are clinically heterogeneous, spanning molecular subtypes and linking to poor prognosis. Reportedly, CNV profiles also influence drug responses; integrating them with mRNA expression may enhance the accuracy of drug sensitivity predictions, supporting advances in personalized therapy.

Given CNV–COAD outcomes associations, we integrated SCNA data for MACC1 with TCGA-COAD RNA-sequencing data. ssGSEA of differentially expressed genes showed downregulated genes enriched in pathways related to ketone body synthesis and degradation. Subsequently, we assessed MACC1’s link to drug sensitivity across multiple cell lines. Sensitivity analysis revealed that high MACC1 levels correlated with responsiveness to several pharmacological agents. Specifically, CEP-710, FK-866, AZD4547, and TW37 exhibited a significant negative correlation, whereas BRD-A86708339 and Lapatinib were positively correlated with high MACC1.

MACC1 is a critical molecular connector linking metabolic syndrome, marked by abdominal obesity, insulin resistance, hyperglycemia, and hypertension, to elevated COAD risk. SCNAs and CNVs are common genomic features in human cancers and drive tumorigenesis by modulating gene expression and underlying molecular mechanisms. In COAD, CNVs are strongly associated with prognosis. Using the LinkedOmics platform, we integrated MACC1 CNV data from the TCGA-COAD cohort (n = 629) with RNA-sequencing profiles. The volcano plot (Fig. 6A) identified genes positively correlated (dark red dots) or negatively correlated (dark green dots) with MACC1-SCNAs. Heatmaps visualized the most significant genes (Fig. 6B, C). Functional enrichment of up-- and down-regulated sets revealed key biological processes associated with MACC1 CNV status (Fig. 6D). Drug sensitivity analyses via GDSC and CTRP databases (Fig. 6E) showed that MACC1 inversely correlated with sensitivity to CEP-710, FK-866, AZD4547, and TW37, but positively correlated with BRD-A86708339 and Lapatinib.

In conclusion, MACC1 regulates COAD progression as a regulatory hub integrating oncogenic signals to drive malignancy and therapy resistance. Current research reveals both challenges and opportunities in targeting MACC1. Advances in structural biology, biomarker refinement, and targeted therapies promise to translate insights into clinical interventions. Ultimately, these efforts will contribute to improved prognostication, personalization, and outcomes for patients with COAD.

## Methods

### Gene of interest filtration

GEPIA2[26] (http://gepia2.cancer-pku.cn) is an interactive web platform designed for gene expression profiling that uses data from 8,587 normal and 9,736 tumor samples available in the GTEx repository[27] (https://gtexportal.org/home/). From the GTEx dataset, 275 COADCOAD cases were extracted, which comprised 275 tumor samples alongside 41 matched adjacent normal tissues. Within this COAD cohort, 390 genes most significantly associated with survival were identified, considering the median expression as the grouping threshold. In addition, applying a significance criterion of adjusted p-value < 0.05 and a |log2 fold change| > 1 yielded 5,222 differentially expressed genes. These two gene sets were subsequently intersected via the online tool at https://www.helixlife.cn/main/, yielding 109 overlapping genes, as illustrated in a Venn diagram.

### Integrated bioinformatics analysis

To perform functional enrichment analysis, the GO[28] and KEGG[29-31] pathway annotations for the 109 candidate genes were analyzed using two independent online platforms. GO term enrichment and annotation were conducted with the Xiantao tool (https://www.helixlife.cn/main/), which supports batch gene annotation and highlights biologically relevant GO terms associated with the input gene set. Moreover, the KEGG pathway mapping and visualization were conducted via the OMICSHARE cloud platform, an open-access tool for functional enrichment analysis and data visualization (http://www.omicshare.com/tools/).

### Investigation of expression

UALCAN (http://ualcan.path.uab.edu) is a publicly accessible web platform designed for the comprehensive analysis of cancer OMICS data [32]. In this study, this tool was employed to examine MACC1 transcription across 33 cancer types based on TCGA data[33]. For a focused analysis on COAD, the TCGA-COAD dataset comprising 521 cases was analyzed to evaluate the MACC1 mRNA expression. Additional information regarding this cohort is available through the TCGA portal (https://portal.gdc.cancer.gov/). Relevant graphs and plots were then generated to visualize the expression profiles. Furthermore, the protein expression of MACC1 in the COAD tissues was investigated with reference to the CPTAC-COAD dataset from the CPTAC (accessible at https://pdc.cancer.gov/pdc/)[34]. This dataset includes 100 cases, with 97 tumor samples paired with 100 adjacent normal tissues. Protein quantification was performed using isobaric labeling-based tissue mass spectrometry (tandem mass tag-10 protocol), as detailed in the corresponding methodology[35]. The expression data were normalized within and across the sample profiles, with the results presented as Z-scores, representing standard deviations from the median. The same CPTAC dataset was used to explore the expression profiles associated with the key signaling pathways, including HIP, mTOR, MYC/MYCN, RTK, SWI-SNF complex, Wnt, p53/Rb-related, chromatin modifier, and NRF2 pathways.

### Survival rate analysis

Kaplan–Meier (KM) survival analysis was performed to assess the relationship between target gene expression and overall survival (OS) in COAD patients. Using the GEPIA2 database[26], the prognostic significance of MACC1 expression was evaluated in a cohort of 270 cases. The patients were stratified into high- and low-expression groups based on the median expression value as the cutoff. Hazard ratios (HRs) along with 95% confidence intervals (CIs) were derived from Cox proportional hazards models, indicated by dotted lines in the plot, and log-rank p-values were calculated to determine statistical significance.

### Immune infiltration analysis

RNA-seq data (in the TPM format) and the corresponding clinical information for the TCGA-COAD project were obtained from the TCGA database via the GDC portal (https://portal.gdc.cancer.gov). These data, originally processed by the STAR pipeline, were subsequently organized and analyzed using R software (version 4.2.1). The ssGSEA algorithm was then applied to quantify the levels of immune cell infiltration within the tumor microenvironment. Spearman’s correlation analysis was performed to assess the associations between immune infiltration scores and gene expression. All data visualization was conducted using the ggplot2 R package (version 3.4.4). This study investigated the relationship between MACC1 expression and immune cell infiltration in COAD. **(A)** Correlation between MACC1 mRNA expression and the abundance of different tumor-infiltrating immune cells. Specific associations are depicted for **(B)** aDC, **(C)** CD8+ T cells, **(D)** cytotoxic cells, **(E)** Tcm (central memory T cells), and **(F)** T helper cells.

### Somatic copy number alteration analysis

LinkedOmics (https://www.linkedomics.org/login.php) is a public data portal that integrates multi-omics datasets across all 32 TCGA cancer types and 10 CPTAC cohorts[36]. To investigate the expression profile of MACC1 and its correlation with clinical outcomes in COAD, we performed an integrated analysis using this platform. We first conducted a joint examination of MACC1 copy number alterations (SCNA) and RNA-seq data from 629 TCGA COAD samples (available via https://pdc.cancer.gov/pdc/). This analysis assessed the association between MACC1 CNVs and their transcript levels, applying thresholds of an adjusted p-value < 0.05 for statistical significance and a log2 fold change > 1 for the magnitude of CNV. The objective was aimed at determining whether CNVs influence the MACC1 expression in COAD as well as to evaluate their prognostic relevance. All procedures followed established bioinformatics protocols, and the results were interpreted using LinkedOmics’ built-in visualization tools.

### Drug sensitivity analysis

The Gene Set Cancer Analysis (GSCA) database (available at GSCA: Gene Set Cancer Analysis (wchscu.cn)) provides a platform for comprehensive cancer-related gene set investigations, including analyses of mRNA expression, mutational profiles, immune infiltration, and drug resistance. This resource aggregates multi-dimensional genomic data from more than 10,000 samples across 33 cancer types in TCGA, along with drug sensitivity information from over 750 small-molecule compounds documented in the Genomics of Drug Sensitivity in Cancer (GDSC) and the Cancer Therapeutics Response Portal (CTRP). By using GSCA, the researchers evaluated drug-sensitivity patterns associated with specific genes, such as *MACC1*, in different cell lines. The platform thus supported the integration of clinical genomic data with pharmacological datasets to help identify the potential therapeutic candidates.

## Data availability

The data utilized in this analysis are publicly accessible from the following repositories: the Genotype-Tissue Expression (GTEx) portal (https://gtexportal.org/home/), TCGA (https://portal.gdc.cancer.gov/), and the CPTAC (https://gdc.cancer.gov/).

## Acknowledgements

Not applicable.

## Author contributions

Conceptualization, Yi Zhang and Peng Sun; methodology and software, Zhenkun Chen; validation and formal analysis, Chun Zheng; investigation, Jie Zhang; resources, Yang Lu; writing—original draft preparation, Zhenkun Chen, Chun Zheng; writing—review and editing, Zhenkun Chen; visualization, Yi Zhang; project administration, Yi Zhang. All authors have read and agreed to the published version of the manuscript.

## Additional information

Correspondence and requests for materials should be addressed to Yi Zhang and Peng Sun.

